# TRPV1-Mediated Delivery of Chloroprocaine, a Local Anesthetic with High pKa, Produces Pain-Selective Anesthesia Without Neurotoxicity

**DOI:** 10.64898/2026.02.24.707701

**Authors:** Kushnir Yishai, Penker Sapir, Lev Shaya, Binshtok M. Alexander

## Abstract

**Background:** Conventional local anesthetics (LAs) provide pain relief but also block motor and non-nociceptive fibers, causing undesirable side effects. Approaches using permanently charged derivatives, such as QX-314, delivered via TRPV1 channels, selectively target nociceptors but are limited by neurotoxicity. We hypothesized that clinically approved LAs with high pKa and low toxicity, such as 2-chloroprocaine (chloroprocaine), which is predominantly protonated and membrane-impermeant at physiological pH, would preferentially enter TRPV1-expressing nociceptors when co-administered with a TRPV1 agonist, thereby producing prolonged pain-selective anesthesia without neurotoxicity.

**Methods:** Whole-cell voltage-clamp recordings from rat dorsal root ganglion (DRG) neurons assessed sodium current inhibition. Behavioral experiments in male Sprague-Dawley rats (N=6 -12/group) evaluated sensitivity to noxious thermal and mechanical stimuli and motor function following intraplantar and perisciatic injections of 0.5% or 2% chloroprocaine combined with capsaicin or cannabidiol (CBD). TRPV1 activation was examined by calcium imaging in hTRPV1-expressing HEK293 cells and in capsaicin- and AITC-sensitive DRG neurons. Neurotoxicity was assessed using propidium iodide uptake, long-term behavioral monitoring (35 days), and immunohistochemistry for TRPV1, ATF-3, and GFAP.

**Results:** Co-application of chloroprocaine with capsaicin enhanced sodium current blockade (76±15%) compared to either drug alone (p<0.05). Perisciatic injection produced thermal analgesia that persisted for 4 hours, whereas motor block resolved within 30 minutes. Unlike other LAs, chloroprocaine did not activate TRPV1 (98% non-responders) and showed no cytotoxicity. Long-term evaluation revealed no hypersensitivity, neuronal loss, or ATF-3 upregulation. Replacing capsaicin with CBD preserved prolonged (∼ 3.5 h) pain-selective anesthesia without motor impairment.

**Conclusions:** TRPV1-mediated delivery of high-pKa LA chloroprocaine produces prolonged, nociceptor-selective anesthesia without neurotoxicity. Combining a clinically approved high-pKa LAs with low intrinsic toxicity and TRPV1 agonists such as CBD offers a potentially safe strategy for pain-selective regional anesthesia, with implications for perioperative and chronic pain management.

## Introduction

Pain has profound effects on healing, well-being, and quality of life, as well as significant economic costs^1^. Local anesthesia is widely used for both acute and chronic pain management^2^. Local anesthetics (LA) act primarily as voltage-gated sodium channel (Na_V_) blockers, preventing conduction along nociceptive pathways and producing effective analgesia. However, LAs also inhibit motor, non-nociceptive, and autonomic fibers, leading to undesirable side effects. Consequently, efforts have focused on developing approaches to selectively block pain pathways by either targeting nociceptor-selective Na_V_s^3–5^, or selectively targeting nociceptor fibers^6,7^. The latter is based on the use of charged LA derivatives, such as QX-314. Because conventional LAs bind to the cytoplasmic side of Na_V_ channels, they must cross the neuronal membrane to block the channels and thereby inhibit conduction^8–10^. Charged LAs penetrate lipid membranes poorly due to their hydrophilic nature and are therefore ineffective when used alone^6,9^. However, they can penetrate via the aqueous pores of cation-nonselective channels, such as the transient receptor potential vanilloid 1 (TRPV1) channel^6,11^. TRPV1 is widely expressed on nociceptive neurons and responds to noxious heat, low pH, and several ligands, including capsaicin^12^. We and others have shown that co-administration of TRPV1 activators facilitates selective entry of QX-314 into nociceptor neurons and produces selective analgesia without affecting motor or non-nociceptive sensory functions^6,11,13–16^. However, this combination is unsuitable for clinical use for several reasons: activation of TRPV1 channels by capsaicin produces prominent pain before QX-314 enters nociceptors to produce analgesia^17^. This problem was addressed by using LA, lidocaine, to activate the TRPV1 channel instead of capsaicin^17^. It has been shown that lidocaine-induced activation of TRPV1 channels is sufficient to allow entry of QX-314 into nociceptive neurons, such that co-application of lidocaine and QX-314 induces a prolonged differential nociceptive block without injection pain^17^. However, the use of QX-314 was also deemed unsuitable due to its neurotoxicity and narrow therapeutic index^18–21^.

LAs are weak bases with pKa values above physiological pH, existing in equilibrium between ionized and non-ionized forms. Only the non-ionized form penetrates the neuronal membrane^22,23^. Thus, molecules with a high pKa, when injected at physiological pH, will be predominantly protonated and less likely to penetrate the cell membrane. We hypothesized that opening large-pore, cation-nonselective channels on nociceptor neurons may provide a “gate” for the protonated fraction of LA to enter these neurons, leading to pain-selective anesthesia. Among clinically used LAs, 2-chloroprocaine (chloroprocaine) has the highest pKa (9.1)^24^. Despite this, it is clinically used for short-acting regional anesthesia via multiple routes^25^ and exhibits efficacy comparable to other agents^26,27^. Importantly, the adverse effects of chloroprocaine are lower than those of other LAs^28^. We hypothesized that, given its high pKa and limited membrane permeability, chloroprocaine, when co-administered with a TRPV1 agonist, may preferentially enter nociceptors through open TRPV1 channels. Thus, combining chloroprocaine, a clinically used LA with a high pKa, wide therapeutic index and low toxicity, with a TRPV1 agonist will provide prolonged, pain-selective analgesia with minimal motor blockade, sensory impairment, or cytotoxicity.

## Materials and Methods

### Animals

All animal procedures were approved by the Ethics Committee of the Hebrew University (MD-18-15551-5, MD-24-17427-5). Male Sprague-Dawley rats weighing 200-250 g were used for all behavioral assessments. For DRG production, rats weighing 100-150 g were used. Males were predominantly used in previous TRPV1-mediated pain-selective anesthesia studies^6,14,18^. Therefore, only males were used to reduce the number of animals in the study^29^.

### Behavioral experiments

#### Animal care

For the behavioral experiments, the rats were housed in the same room where they were tested for at least a week before the experiment. Temperature and humidity were controlled and monitored under a 12-hr. light/dark cycle. All experiments were performed during the light phase. Animals were habituated to the handler and testing apparatus for 7 days before the experiment; during habituation, no noxious stimuli were applied.

Intraplantar injections of 10 µL of the test substances or vehicle were performed on the plantar aspect of the left hind paw while the animal was manually restrained. For perisciatic injections, rats received 100 µg/kg medetomidine (Domitor, Orion Pharma, Finland) subcutaneously, and 10 minutes were allowed for the onset of sedation. Rats were then restrained in lateral recumbency, and landmarks (greater trochanter and ischial tuberosity) of the right hind limb were localized. Rats were injected with 100 µL of the test substance or vehicle in immediate proximity to the sciatic nerve using a 27-gauge hypodermic needle attached to a tuberculin syringe, as described previously^17^. To examine the effect of the time interval between injections, the two injections were performed using the same technique, with a variable time interval between injections as required for each drug combination. Unless stated otherwise, chloroprocaine (Chloroprocaine hydrochloride, USP, Rockville, MD) or saline (sodium chloride 0.9%, B Braun, Germany) control was injected first, followed by the TRPV1 agonist (e.g., capsaicin) or vehicle control. Immediately after the last perisciatic injection, rats were administered 500 µg/kg atipamezole (Antisedan, Orion Pharma, Finland) to antagonize the sedative effects of medetomidine. To evaluate the efficacy and accuracy of the sciatic nerve block technique, indicated by complete paralysis of the hind limb, a positive control group of lidocaine 2% (B Braun, Germany) or chloroprocaine 2% was used. T0 was defined as the time of the 2^nd^ (TRPV1 agonist) injection.

#### Preparation of Test Solutions for In Vivo Experiments

Capsaicin (Sigma-Aldrich, St Louis, MO) was dissolved in ethanol to prepare a 5 mg/ml stock, which was then mixed with 5% Tween 20 (Sigma-Aldrich, France) in saline at a ratio of 1:9 to produce a solution with a final capsaicin concentration of 0.5 µg/µl. chloroprocaine was dissolved in saline to achieve the required concentrations. CBD (plant extract, kindly provided by the lab of Dr. Raphael Mechoulam, School of Pharmacy, Hebrew University of Jerusalem, Hadassah Ein Kerem) was dissolved in ethanol to produce a 100 mg/ml stock. The stock was mixed with 5% Tween 20 in saline at a 1:9 ratio to produce a final CBD concentration of 10 µg/µL. Vehicle control for capsaicin and CBD was prepared by mixing ethanol with 5% Tween 20 in saline at a ratio of 1:9. Saline was used as a control for chloroprocaine and lidocaine.

#### Behavioral tests

Sensitivity to a noxious heat stimulus was assessed using the Hargreaves paw withdrawal latency test. After habituation to the handler and examination environment, rats were placed on a plexiglass surface, and a laser beam was directed at the plantar aspect of the hind paw (Ugo-Basile, Varese, Italy). The laser heated the paw, eliciting a behavioral withdrawal response, such as moving, flinching, or licking the paw. The time from initiation of the heat stimulus until the rat withdrew its paw was automatically measured as the paw withdrawal latency (PWL). A 20-second cutoff was set to avoid tissue damage.

Sensitivity to mechanical stimuli was measured using an electronic Von Frey apparatus (Ugo-Basile, Varese, Italy). After habituation, the rats were placed on a mesh platform. A blunt-tipped needle was placed on the plantar aspect of the hind paw, and pressure was increased gradually at the indicated rate until the paw was withdrawn. The pressure (in Newtons, N) required to elicit a withdrawal reflex was recorded as the mechanical pain threshold.

Motor function was assessed using a climbing task, as described in Binshtok et al. 2007^6^. Rats were placed on a vertical grid, and hind-limb use was scored 0 (normal function), 1 (impaired motor function, e.g., weakness or failure to grasp), or 2 (complete paralysis).

All behavior tests were performed by a single assessor (YK), blinded to treatment allocation.

### Acutely dissociated dorsal root ganglia (DRG) neurons preparation

#### DRG excision

Rats were deeply anesthetized with isoflurane (Isoflurane, USP Terrell, Piramal Critical Care, Bethlehem, PA, USA) and sacrificed by decapitation. DRG were isolated and dissociated as described previously^30^. Briefly, lumbar DRG neurons (L1-L6) were removed and placed in Dulbecco’s modified Eagle medium with 1% penicillin–streptomycin, then digested with 5 mg/mL collagenase, 1 mg/mL Dispase II (Roche), and 0.25% trypsin, followed by the addition of 0.25% trypsin inhibitor. Cells were triturated with Pasteur pipettes in the presence of DNase I (250 U) and centrifuged through 15% BSA. The cell pellet was resuspended in 1 mL of Neurobasal medium containing B27 supplement (Invitrogen, Carlsbad, CA), penicillin and streptomycin, 10 mM AraC, 2.5S NGF (100 ng/mL, Promega), and glial cell-derived neurotrophic factor (2 ng/mL). Cells were plated onto poly-D-lysine (100 g/mL) and laminin (1 mg/mL)-coated 35-mm tissue culture dishes. Cells were kept in the incubator at 37°C with 5% carbon dioxide.

#### Expression of hTRPV1

HEK^293^ cells with stable expression of human TRPV1 (hTRPV1, kindly provided by the lab of Dr. Baruch Minke, Department of Medical Neurobiology, Hebrew University, Jerusalem) were cultured and prepared as previously described^20^ in standard DMEM (Biological Industries) with 10% FBS (Biological Industries), 100 U/ml penicillin and 100 μg/ml streptomycin (Biological Industries), and 5% L-glutamine (Biological Industries). 5 μg/ml blasticidin (InvivoGen, Toulouse, France) and 0.35% Zeocin (InvivoGen, Toulouse, France) were added to maintain stable hTRPV1 expression. For tetracycline induction, 0.1 μg/ml (InvivoGen, Toulouse, France) was added to the medium 4-8 h prior to experiments.

#### Ca^2+^ imaging

Fluorescent Ca^2+^ imaging was performed using an inverted microscope (Nikon Eclipse Ti) equipped with an Epi-Fl attachment, a perfect focus system (Nikon, Tokyo, Japan), and an Exi Aqua monochromator (QImaging). Acutely dissociated DRG neurons or HEK^293^ cells expressing hTRPV1 were loaded with fura-2 acetoxymethyl (fura-2 AM, stock in DMSO) for 45 to 60 minutes in a standard extracellular bath solution (SES) composed of (in mM): 145 NaCl, 5 KCl, 2 CaCl_2_, 1 MgCl_2_, 10 glucose, and 10 HEPES. They were then rinsed for 45 to 60 minutes to de-esterify intracellular AM esters. Intracellular [Ca^2+^] was measured fluorometrically as the ratio of excitation at 340 nm and 380 nm (F = ΔF340/380). Emission was collected at 510 nm. At the beginning of the experiment, cells were bathed in SES. After 5 minutes, cells were perfused with SES composed of (in mM): 145 NaCl, 5 KCl, 1 MgCl_2_, 2 CaCl_2_, 10 glucose, and 10 HEPES. Under these conditions, the substance under examination (e.g., chloroprocaine, capsaicin, AITC) was perfused into the test plate at varying concentrations according to the study’s requirements. SES was perfused following drug administration to ensure washout. In all experiments, images were acquired at 3-second intervals, monitored online, and analyzed offline using Nikon Elements AR Software (Nikon). For quantification of changes in fluorescence intensity, values of fluorescence at specific time points (F) were normalized to the baseline signal (F - F_0_).

#### Electrophysiological recordings

Recordings were performed from small (∼25 μm) dissociated rat DRG neurons, up to 48 hours after plating. In some experiments, 1 µM capsaicin was added to the bath solution at the end of the experiment, and the resulting inward current was measured. Cell diameter was measured online using Nikon Elements AR software (Nikon), from images acquired using a CCD camera (Q-Imaging). Whole-cell membrane currents were recorded using a voltage-clamp mode, with a MultiClamp 700B amplifier (Molecular Devices), at RT (24 ± 2°^C^). Data were sampled at 20 kHz and were low-pass filtered at 10 KHz (−3 dB, 8-pole Bessel filter). Patch pipettes (2-5 MΩ) were pulled from borosilicate glass capillaries (1.5/1.1mm OD/ID; Sutter Instrument Co, Novato, CA) on a P-1000 puller (Sutter Instrument Co., Novato, CA) and fire-polished (LW Scientific, Lawrenceville, GA). Command voltage and current protocols were generated with a Digidata 1440A A/D interface (Molecular Devices, San Jose, CA). Data were digitized using pCLAMP 10.3 (Molecular Devices). Data averaging and peak detection were performed using Clampfit 10.3 software. Data were fitted and analyzed using Origin 9 (OriginLab) and Matlab.

The extracellular solution contained (in mM): 145 NaCl, 5 KCl, 1 MgCl_2_, 10 HEPES, and 10 D-glucose. The pH was adjusted to 7.4. Pipette potential was zeroed before seal formation. Intracellular solution for voltage clamp recordings contained the following (in mM): CsCl, 110; CsOH, 25; NaCl 10, MgCl_2_, 2; CaCl_2_ 1; EGTA 11; HEPES, 10 (pH = 7.4 with CsOH). Peak sodium current amplitudes were measured following 150 ms depolarizing steps to a range of test potentials in 10 mV increments, from a holding potential of −70 mV. Currents were assessed before and after application of the examined substance, e.g. chloroprocaine with or without capsaicin at varying concentrations. Recordings were obtained from multiple animals (n=2-4 rats) to minimize pseudoreplication.

#### Cell viability assays

Cell viability was assessed by Propidium Iodide (PI) staining according to the manufacturer’s protocol (FITC Annexin V Apoptosis Detection Kit 1; BD Pharmingen, Germany) and as described previously^21^. Cells were cultured in 35 mm tissue culture dishes and, upon reaching 80% confluence, exposed to the tested substances for 1, 3, 6, 12, and 24 hours. Staining was performed by adding 5 μl PI and incubating for 15 min at room temperature in the dark. Specimens were analyzed using light and fluorescent microscopy, and the percentage of non-viable cells (non-viable cells / total cell count per field) was calculated manually.

#### Immunohistochemistry

Rats were deeply anesthetized with isoflurane and fixed by intracardiac perfusion with 100 mL of 4% paraformaldehyde (PFA) freshly prepared in phosphate-buffered saline (PBS, pH 7.4) at 4°C. The dorsal root ganglia of lumbar segments L4-L6 was then immediately dissected out, followed by immersion for 1 hour in 4%PFA at 4°C, and cryoprotected by overnight immersion in 30% sucrose in PBS at 4°C. Tissue samples were then frozen in optimal cutting temperature (OCT) medium, and 15-µm cryosections were collected onto Superfrost Plus slides and stored at 220°C.

Slides were first thawed at room temperature (RT), washed in PBS, and incubated for 7 minutes in a permeabilization solution (0.5% Tween-20 and 1% Triton X-100 in PBS), followed by incubation of 1 hour with blocking solution (0.3% Triton X-100, and 5% donkey serum in tris-buffered saline [TBS]). Sections were then incubated overnight at 4°C with primary antibodies, diluted in antibody solution (0.1% Tween-20 and 3% bovine serum albumin in PBS). Then the samples were washed 3 times for 10 minutes in PBS, followed by incubation in the dark with fluorescent secondary antibodies, diluted in antibody solution (0.1% Tween-20 and 3% bovine serum albumin in PBS) for 1 hour at room temperature. Finally, sections were washed 3 times for 10 minutes in PBS, then dried out, and mounted with Vectashield with DAPI and coverslipped.

#### Antibodies

The triple labelling was performed using (1) mouse monoclonal anti rat ATF-3 (1:500, Santa Cruz, Cat# sc-518032) with Alexa Fluor 647-conjugated AffiniPure Fab Fragment Goat Anti-Mouse (1:500, Jackson Immunoresearch, Cat# 115-607-186) as secondary antibody; (2) chicken Anti-rat GFAP (1:1000, Millipore, Cat# AB5541) with Alexa Fluor 488-conjugated AffiniPure Donkey Anti-chicken (1:1000, Jackson Immunoresearch, Cat# 706-545-155) secondary antibody and (3) rabbit anti-ratTRPV1 (1:1000, Alomone labs, Israel, Cat# ACC-030) with Alexa Fluor 555-conjugated IgG H&L Goat Anti-Rabbit (1:500, Abcam, Cat# AB-ab-150078) secondary antibody.

#### Image analysis for immunohistochemistry

Images were acquired using a confocal microscope (Nikon Ti2E) equipped with a 20× objective. Images were processed using NIH ImageJ software (Bethesda, MD, USA), and analyses were performed on maximum intensity projections. For quantification of GFAP immunofluorescence, a segmented line was traced along the borders of the dorsal root ganglion (DRG), and a background region was selected within the same image. Mean fluorescence intensity was measured using the ImageJ “Measure” function. Background intensity was subtracted from tissue intensity to obtain the average GFAP fluorescence intensity. Total cell counts were determined by quantifying DAPI-stained nuclei. The percentages of TRPV- and ATF-positive cells per section were calculated as (number of positively stained cells / number of DAPI-stained nuclei) × 100.

#### Statistical Analysis

The experimental outcomes of responsiveness to noxious thermal or mechanical stimuli, sodium current amplitudes, and cell viability assays generated continuous outcome data suitable for parametric and nonparametric statistical analyses. Group comparisons were performed using Student’s *t*-test or one- or two-way analysis of variance (ANOVA) for normally distributed data. When applicable, Bonferroni or Tukey adjustments were used to account for multiple comparisons. For non-normally distributed data, group differences were analyzed using the Kruskal-Wallis test followed by the two-stage linear step-up procedure of Benjamini, Krieger, and Yekutieli.

Motor activity was scored on an ordinal scale (0 = no block, 1 = partial block, 2 = complete block) at baseline and multiple time points following injection. Because the outcome variable was ordinal and repeatedly measured within animals, data were analyzed using a generalized estimating equations (GEE) with an ordinal cumulative logit link function. Treatment group, time, and their interaction were included as fixed effects, and animal identity was treated as a repeated factor with an exchangeable working correlation structure. This model was used to assess overall group effects, time effects, and group × time interactions. To further characterize differences at individual time points, between-group comparisons were performed using Mann–Whitney U tests with Holm-Bonferroni correction for multiple comparisons. Within each group, changes from baseline were assessed using paired Wilcoxon signed-rank tests comparing baseline with each post-injection time point, with Holm correction applied across time points. In addition, a summary measure of motor block over time was calculated for each animal as the area under the curve (AUC; score*h) using the trapezoidal rule after converting time to hours; AUC values were compared between groups using Mann–Whitney U tests.

For behavioral experiments, an integrative analysis was performed by calculating baseline-subtracted (measured at point 0) area under the curve (AUC) for each animal using trapezoidal integration (0 - 360 min for thermal sensitivity and 0 - 240 min for mechanical sensitivity), with results expressed as seconds*hours or grams*hours, respectively. Individual AUC values were compared across groups using a one-way analysis of variance followed by Tukey’s post hoc test.

ATF-3 and TRPV1 immunoreactivity was quantified as the percentage of ATF or TRPV1-positive neurons relative to the total number of DRG neurons within each analyzed section. For GFAP immunoreactivity mean GFAP-induced fluorescence intensity of each section was calculated. Each data point corresponds to a single histological section. Because sections from different animals were pooled prior to analysis, individual animal identity could not be retained and therefore could not be modeled as a random effect. Consequently, statistical analyses were performed at the section level. All statistical inferences, therefore, reflect variability between sections rather than between animals. Data were analyzed using a two-way linear model with “Treatment” (saline+vehicle, chloroprocaine+vehicle, saline+capsaicin, chloroprocaine+capsaicin) and “Side” (ipsilateral vs contralateral) as fixed factors, including their interaction term. Model assumptions (normality of residuals and homoscedasticity) were evaluated by inspection of residual plots and compared using the Kolmogorov-Smirnov test. When appropriate, post hoc comparisons were conducted using Tukey’s test to correct for multiple comparisons across all Treatment × Side groups.

All statistical tests were two-tailed. Statistical significance was set at *p* < 0.05 after correction for multiple comparisons. Data are presented as mean ± SEM unless otherwise indicated. Analyses were performed using Python and GraphPad Prism.

## Results

### TRPV1 activation enhances chloroprocaine-induced sodium current blockade in nociceptor neurons

Activation of TRPV1 channels provides a large-pore pathway that allows otherwise membrane-impermeant or poorly permeant charged molecules, including quaternary local anesthetic derivatives, to enter nociceptive neurons selectively^11,31–33^. Building on this concept, we hypothesized that opening TRPV1 channels would similarly facilitate intracellular access of the large protonated, charged fraction of high-pKa LA chloroprocaine, thereby enhancing its ability to inhibit voltage-gated sodium currents in nociceptors. If this is true, co-application of chloroprocaine with a TRPV1 agonist such as capsaicin should produce a greater reduction in sodium currents than either agent alone.

To test this hypothesis, we performed whole-cell voltage-clamp recordings of total sodium currents from small-diameter (less than 25 μm) capsaicin-sensitive rat nociceptor DRG neurons. Sodium currents were evoked by 150–ms depolarizing steps from a holding potential of −70 mV, and peak currents were quantified before drug application, after exposure to 100 μM (0.5%) chloroprocaine alone or 1 μM capsaicin alone, and after sequential application of chloroprocaine followed by capsaicin, with a subsequent washout. We used a relatively low concentration of chloroprocaine to achieve about 50% inhibition of sodium current amplitude (**Figure 1**, *blue traces*). We found that co-application of chloroprocaine followed by capsaicin produced a substantial inhibition (76% ± 15%) of peak sodium currents that was significantly stronger than either chloroprocaine (52% ± 17% inhibition) or capsaicin alone (31% ± 18% inhibition **Figure 1**, *middle*). After 5 min washout, in the chloroprocaine+capsaicin group, sodium currents partially recovered but remained reduced by ∼35% compared with baseline, whereas the chloroprocaine -only and capsaicin-only groups showed smaller residual reductions of ∼23% relative to baseline. These findings support the concept that TRPV1 activation enhances chloroprocaine entry into nociceptors, leading to more pronounced and sustained sodium channel blockade.

**Figure 1.**
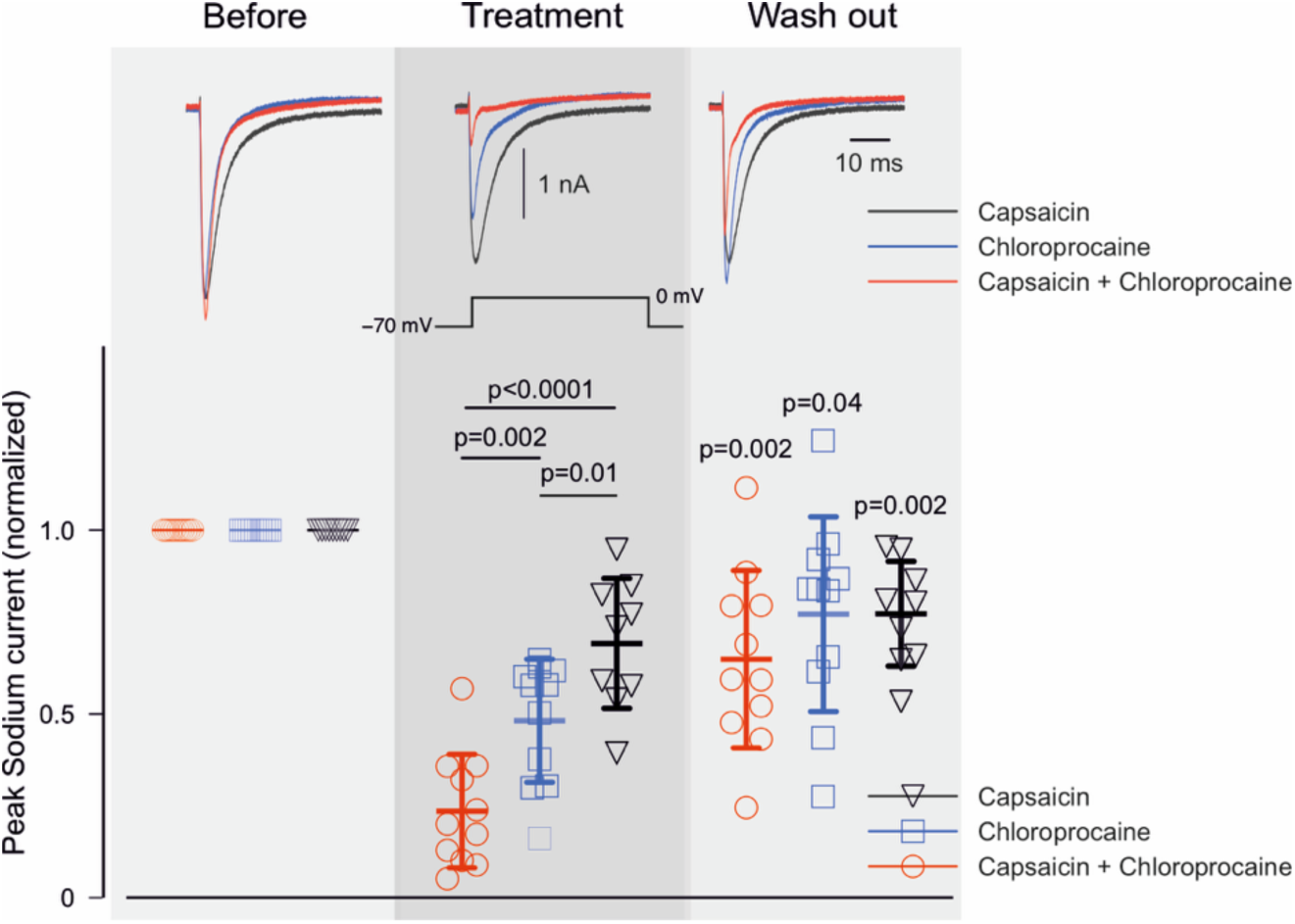
TRPV1 activation enhances chloroprocaine-induced block of the sodium currents in nociceptor neurons. *Upper*, Representative traces of sodium currents recorded from small-diameter rat DRG neurons (<25 μm) before, during treatment (10 min exposure to 100 μM (0.5%) 2-chloroprocaine alone (*blue*), 1 μM capsaicin alone (*black*), or 100 μM chloroprocaine together with 1 μM capsaicin (*red*)), and after washout. Sodium currents were evoked by 150–ms depolarizing steps from a holding potential of −70 mV to test potentials between −60 and 0 mV. *Lower*, Group data summarize normalized peak sodium currents (mean ± SEM) before treatment, during treatment, and after washout. Two-way repeated-measures ANOVA followed by Tukey’s post hoc test; exact p-values are shown on the graph. Each symbol represents one neuron; n = 11, 9, and 11 neurons from N=4, 2, and 3 rats for chloroprocaine alone, capsaicin alone, and chloroprocaine with capsaicin, respectively.

### Capsaicin facilitates the analgesic effect of chloroprocaine

We next examined whether enhanced sodium current block translates into nociceptor-selective analgesia or differential block *in vivo.* We first examined whether adding capsaicin is sufficient to facilitate chloroprocaine-induced analgesia. We used a low dose of chloroprocaine (0.5%), which, when injected intraplantar followed 2 minutes later by the injection of vehicle for capsaicin (chloroprocaine+vehicle, *see Methods*), did not affect the latency of response to a noxious thermal stimulus, measured as the time to withdrawal from a standardized radiant noxious heat stimulus applied to the plantar surface of the hind paw (**Figure 2A**, *blue squares*). Injections of saline followed by the 0.05% capsaicin, followed by the injection of saline (saline+capsaicin) led to a significant but brief (∼30 min) decrease in thermal latency (**Figure 2A**, *black inverted triangles*).

**Figure 2.**
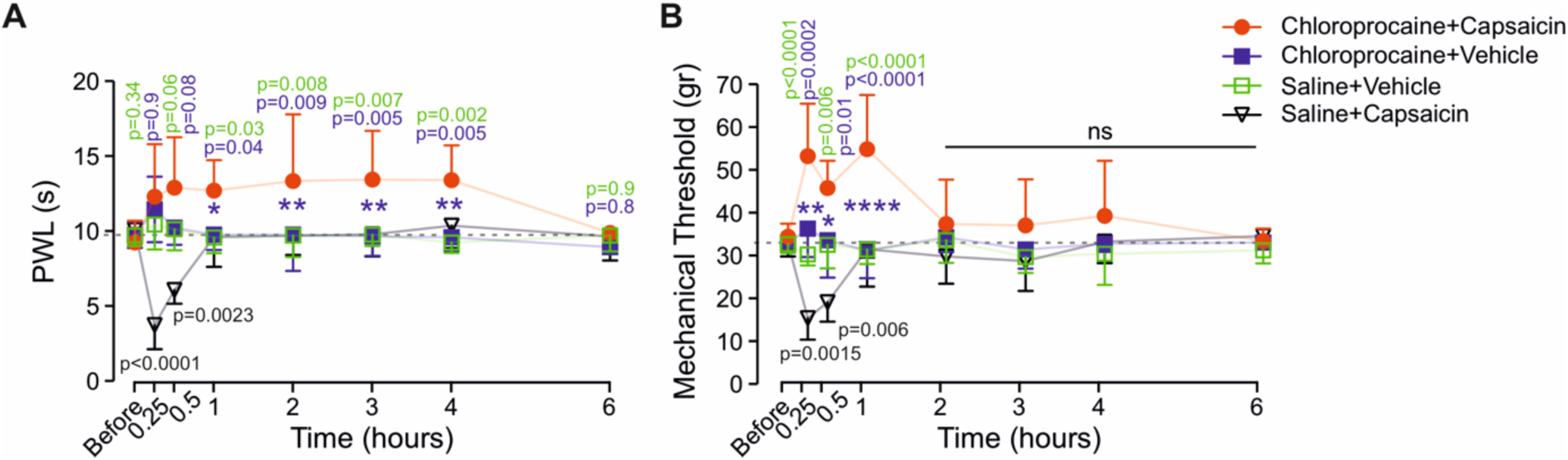
Capsaicin facilitates chloroprocaine-induced analgesia after intraplantar injection. (**A**) Time course of changes in sensitivity to noxious thermal stimuli, measured as thermal paw-withdrawal latency (PWL) following intraplantar injection of 0.5% chloroprocaine followed by 0.05% (0.5 µg/µl) capsaicin (chloroprocaine+capsaicin), 0.5% chloroprocaine followed by vehicle (chloroprocaine+vehicle), saline followed by 0.05% capsaicin (saline+capsaicin), or saline followed by vehicle (saline+vehicle). (**B**) Time course of changes in mechanical withdrawal thresholds after the same treatments as in *A*. Data are presented as mean ± SEM. Two-way repeated-measures ANOVA followed by Tukey’s multiple-comparison test at individual time points. Statistical comparisons between the chloroprocaine+capsaicin and chloroprocaine+vehicle groups are indicated in blue, and between chloroprocaine+capsaicin and saline+vehicle in green. Exact p-values are shown on the graph. The dotted lines indicate the mean value before treatment. N = 6 rats per group.

A combination of 0.5% chloroprocaine and 0.05% capsaicin (chloroprocaine+capsaicin), injected consequently, into the plantar surface of the hind paw, produced a prolonged increase in thermal latency that was significantly longer than chloroprocaine+vehicle (**Figure 2A**, AUC_Chloroprocaine+Capsaicin_ = 19.9 ± 2.2 vs. AUC_Chloroprocaine+Vehicle_ = 0.14 ± 3.8, s*h, n = 6 rats, p = 0.002). The significant increase in response latency began 60 min after injection and persisted for 6 hours (**Figure 2A**, *red circles*).

Similarly, injection of 2-chloroprocain+capsaicin increased the mechanical threshold for eliciting a flexion reflex, as measured by the von Frey apparatus applied to the plantar skin (**Figure 2B**). The increase began 15 min after injection and lasted for 1 hour. Injection of 0.5% chloroprocaine with vehicle instead of capsaicin did not produce any significant change in mechanical sensitivity (**Figure 2B**, AUC_Chloroprocaine+Capsaicin_ = 98.1 ± 9.9 vs. AUC_Chloroprocaine+Vehicle_ = 5.1 ± 5.2, g*h, n = 6 rats, p < 0.001; see also **Figure 2A**). These data suggest that activation of TRPV1 channels permits the entry of the protonated fraction of chloroprocaine into nociceptor neurons, thereby facilitating its analgesic effect.

### TRPV1-targeted perisciatic delivery of chloroprocaine produces prolonged pain-selective anesthesia

If, as our results suggest, the enhanced effect of chloroprocaine when applied with capsaicin is due to facilitated entry of chloroprocaine via capsaicin-induced opening of TRPV1 channels, the effect should be prominent only in TRPV1-expressing nociceptor neurons and should not affect other modalities. To test whether a chloroprocaine+capsaicin combination can induce a long-lasting, nociceptor-specific block, we applied this combination to a major nerve that also contains motor fibers. We administered perisciatic injections of 0.5% chloroprocaine, followed by capsaicin or its vehicle at a fixed 10-minute interval, with the idea that the protonated form of chloroprocaine would be extracellularly available to enter TRPV1 channels upon activation by capsaicin^6^. We assessed thermal and mechanical withdrawal thresholds and compared them with motor performance on a vertical grid over time (*see Methods*). We showed that perisciatic co-administration of saline followed by capsaicin (saline+capsaicin group) did not affect motor behavior (**Figure 3A**). The application of 0.5% chloroprocaine, followed by either capsaicin or vehicle, produced a similar impairment in motor function that began at 15 minutes and resolved after 30 min following injection, compared with the group injected with saline followed by vehicle for capsaicin (**Figure 3A**, AUC_Chloroprocaine+Capsaicin_ = 1.1 ± 0.8 vs. AUC_Chloroprocaine+Vehicle_ = 0.9 ± 0.9, n = 12 rats, p = 0.3).

**Figure 3.**
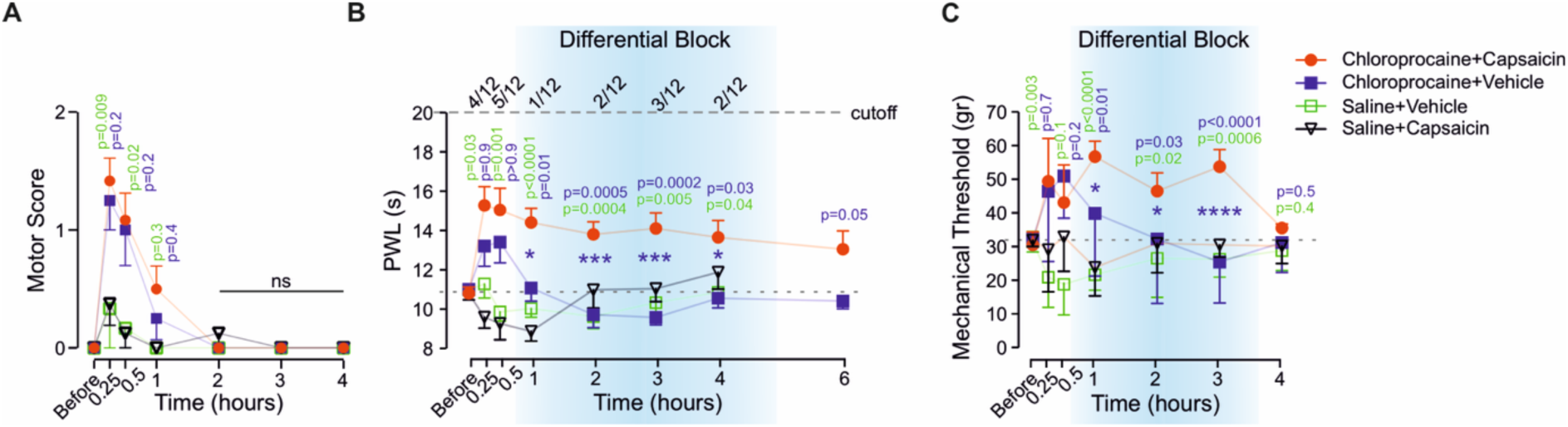
Perisciatic TRPV1-targeted delivery of chloroprocaine produces prolonged pain-selective block. **(A)** Changes in motor scores (0-2) over time after perisciatic injection of 0.5% chloroprocaine followed by 0.05% (0.5 µg/µl) capsaicin (chloroprocaine+capsaicin), 0.5% chloroprocaine followed by vehicle (chloroprocaine+vehicle), saline followed by 0.05% capsaicin (saline+capsaicin), or saline followed by vehicle (saline+vehicle). (**B+C**) Changes in the sensitivity to noxious thermal stimuli (paw withdrawal latency, PWL, **B**) and mechanical withdrawal thresholds (**C**) over time following the same treatments as in *A*. Shaded regions indicate the period of differential block, during which nociceptive function remained suppressed after the motor block resolved. The dotted lines indicate the mean value before treatment. The dashed lines indicate the cutoff, and the numbers above indicate the number of animals that reached the cutoff. Data are mean ± SEM. Motor scores were analyzed using generalized estimating equations; between-group differences at individual time points were further assessed with Mann-Whitney tests with Holm-Bonferroni correction. Thermal and mechanical thresholds were analyzed with two-way repeated-measures ANOVA followed by Tukey’s multiple-comparisons test. N = 12 rats per group for motor and thermal measures; n = 6 rats per group for mechanical thresholds. Statistical comparisons between the chloroprocaine+capsaicin and chloroprocaine+vehicle groups are indicated in blue, and between chloroprocaine+capsaicin and saline+vehicle in green. Exact p-values are shown on the graph.

Notably, perisciatic injection of chloroprocaine followed by capsaicin produced a robust, long-lasting decrease in sensitivity to a noxious thermal stimulus (**Figure 3B**). The increase in thermal paw-withdrawal latency began 15 minutes after treatment and persisted for at least 4 hours, outlasting motor block and producing a differential block of over three hours (**Figure 3B**). Importantly, treatment with chloroprocaine+vehicle produced a significantly shorter increase in thermal latency (AUC_Chloroprocaine+Capsaicin_ = 13.6 ± 3.4 vs. AUC_Chloroprocaine+Vehicle_ = 0.6 ± 1.5, s*h, n = 12 rats, p = 0.003), lasting only 30 min, similar in duration to the motor block (**Figure 3A, B**). As expected, application of saline followed by capsaicin produced a short-lasting (15 min) increase in sensitivity to noxious thermal stimuli (**Figure 3B**). The increase in mechanical withdrawal thresholds following treatment with chloroprocaine+capsaicin began 15 min after treatment and lasted for 3 hours post-injection (**Figure 3C**), whereas the reduction in responsiveness following injection of chloroprocaine+vehicle lasted 1 hour (**Figure 3B, C**, AUC_Chloroprocaine+Capsaicin_ = 51. 2± 15.1 vs. AUC_Chloroprocaine+Vehicle_ = 9 ± 11.5, g*h, n = 6 rats, p = 0.02).

Collectively, these data show that TRPV1-facilitated delivery of chloroprocaine produces a differential block in which nociceptive function remains suppressed hours after motor function has recovered.

Interestingly, increasing the chloroprocaine concentration to 2% did not improve the differential block. It prolonged motor impairment for 1 hour (**Figure S1A**) and decreased sensitivity to a noxious thermal stimulus for 2 hours (compared with the saline+vehicle group; **Figure S1A**).

### Chloroprocaine–capsaicin combinations are non-neurotoxic and do not produce long-term sensory deficits

Because QX-314-based TRPV1-mediated approaches are neurotoxic^18^, we next evaluated whether substituting QX-314 with clinically approved chloroprocaine would maintain safety at the cellular and tissue levels.

We and others have previously demonstrated that QX-314-induced toxicity is mediated by TRPV1 channel activation^20,21^. Therefore, we first examined whether chloroprocaine also activates TRPV1 channels. We expressed human TRPV1 channels (hTRPV1) in HEK293 cells and monitored changes in intracellular calcium following the application of chloroprocaine and capsaicin. The absolute majority of capsaicin-sensitive HEK293-hTRPV1 cells did not respond to 1 mM chloroprocaine (98%, 190 out of 194 cells in 4 plates; **Figure 4A**). We then exposed hTRPV1-expressing HEK293 cells to a supraclinical concentration of 100 mM chloroprocaine (3%). Under these conditions, too, the majority of HEK293-hTRPV1 cells did not respond to this high concentration of chloroprocaine (96%, 390 nonresponders out of 410 cells, 4 plates, **Figure S2**).

**Figure 4.**
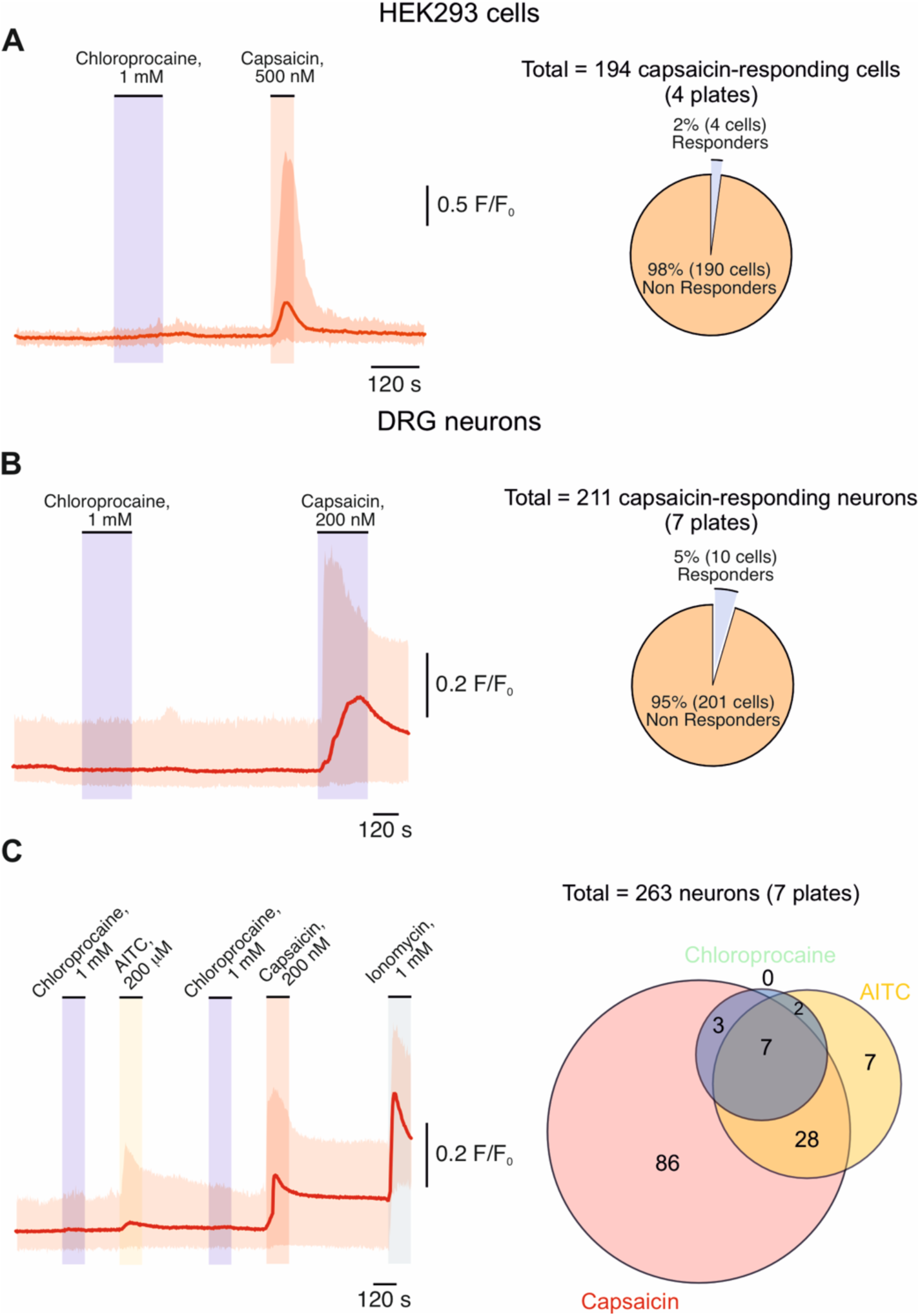
Chloroprocaine does not activate TRPV1 or TRPA1 channels. **(A)** Representative trace of mean changes in Ca^2+^ responses (measured as ratiometric changes in Fura-2 fluorescence; the cloud shows the full range of responses) in HEK293 cells expressing hTRPV1 following application of 1 mM chloroprocaine and 500 nM capsaicin. The pie chart shows the proportion of capsaicin-responsive cells that responded to chloroprocaine. Representative of 4 out of 4 plates. (**B**) Same as *A* but showing changes in Ca^2+^ in capsaicin-sensitive rat DRG neurons during application of 1 mM chloroprocaine and 200 nM capsaicin, with corresponding proportion of responders. Representative of 7 out of 7 plates. (**C**) Same as *B* but showing the Ca^2+^ responses of DRG neurons expressing TRPA1 and TRPV1 channels to chloroprocaine, AITC, and capsaicin. Representative of 7 out of 7 plates. A Venn diagram summarizes overlap among chloroprocaine-, AITC-, and capsaicin-responsive neurons.

Similarly, application of chloroprocaine did not activate acutely dissociated capsaicin-sensitive (i.e., TRPV1-expressing) rat DRG neurons (201 nonresponders out of 211, 7 plates, N=3 rats, **Figure 4B**). We also show that application of chloroprocaine does not activate AITC and capsaicin-sensitive (TRPA1 and TRPV1 expressing neurons, **Figure 4C**).

Altogether, these data show that, unlike other clinically used LAs, chloroprocaine did not activate TRPV1 or TRPA1 channels. Considering that at least in part the toxicity of QX-314 is mediated via TRP channels activation^21^, our results suggest that chloroprocaine has potentially low toxicity. Indeed, 24-hour treatment with 1 mM chloroprocaine did not cause cell death in HEK293 cells expressing hTRPV1, as assessed by propidium iodide uptake (**Figure 5A**). By comparison, exposure to 1 μM capsaicin under the same conditions produced substantial cell death, evident 12 hours after exposure (**Figure 5A**).

**Figure 5.**
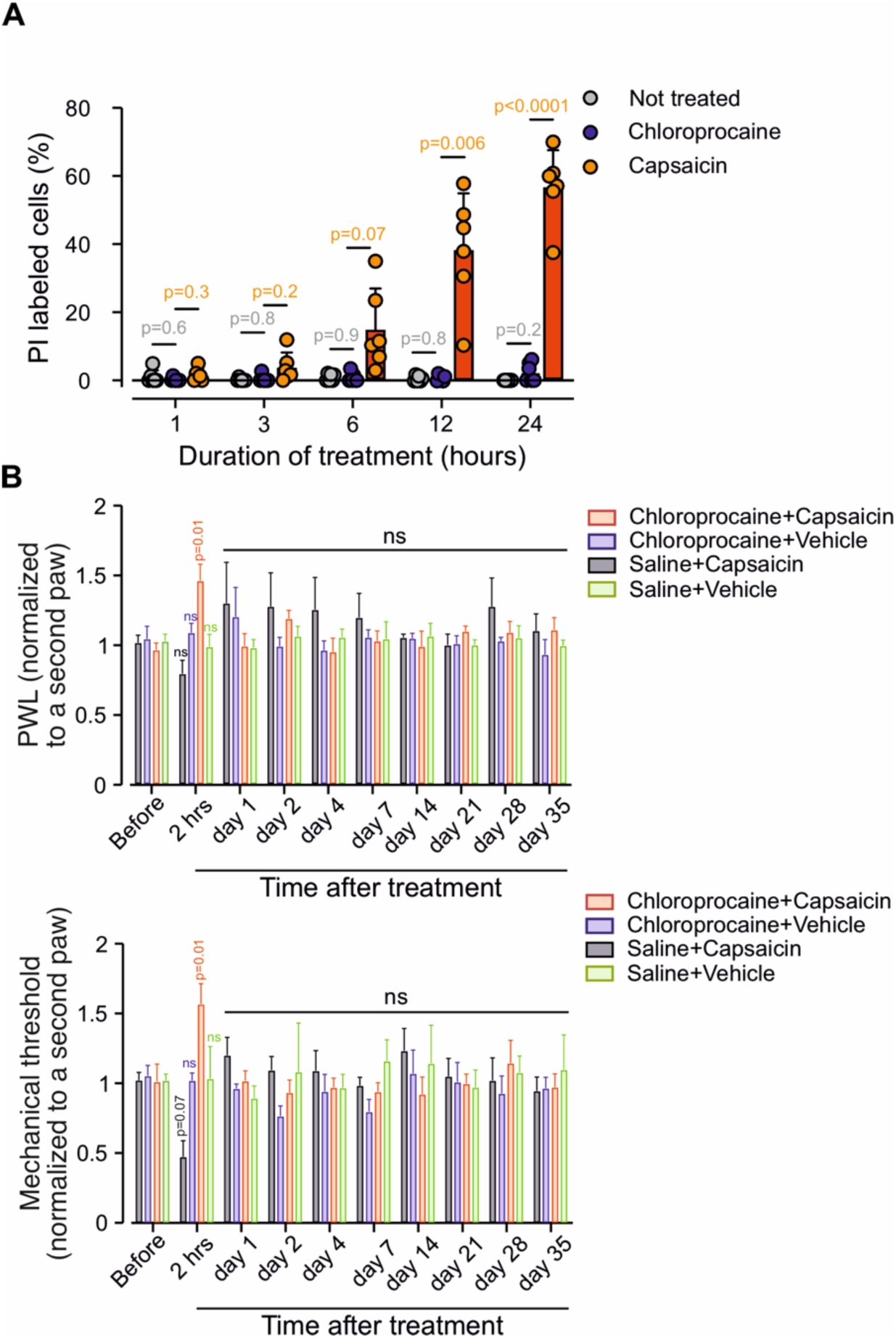
Chloroprocaine-capsaicin combinations are not cytotoxic and do not induce long-term behavioral changes. **(A)** Percentage of propidium iodide-positive HEK293-hTRPV1 cells after exposure to 1 mM chloroprocaine, 1 μM capsaicin, or no treatment for 1-24 h (mean ± SEM). Group differences at each time point were assessed using one-way ANOVA followed by Tukey’s post hoc test. Each point represents an analysis of a single field of view (*see Methods*). (**B**) Long-term changes in the sensitivity to noxious thermal stimuli (PWL, *top*) and mechanical thresholds (von Frey, *bottom*) normalized to the contralateral paw up to 35 days after perisciatic injection of chloroprocaine+capsaicin, chloroprocaine+vehicle, saline+capsaicin, or saline+vehicle (mean ± SEM). Repeated measures over time were analyzed using two-way repeated-measures ANOVA followed by Tukey’s multiple-comparisons test; no significant between-group differences were detected beyond the 2-h time point. N = 6 rats per group.

We next examined whether co-application of chloroprocaine with capsaicin produces long-term neurotoxicity *in vivo*, as observed with QX-314, which induced long-term mechanical hypersensitivity^18^. We assessed long-term changes in thermal and mechanical sensitivity 35 days after perisciatic application of chloroprocaine and capsaicin. As expected from our results shown above (see **Figure 3**), the injection of chloroprocaine+capsaicin elevated thermal and mechanical thresholds 2 hours after injection. However, no differences were detected at later time points up to 35 days, indicating that the analgesic effect was robust yet short-lasting and reversible (**Figure 5B**).

To determine whether perisciatic application of chloroprocaine+capsaicin leads to subclinical neuronal injury, we performed immunohistochemical analysis of DRG harvested 35 days after injection, assessing differences in the number of TRPV1-expressing neurons and in markers associated with neurotoxicity, such as ATF-3 and a glial marker, GFAP^18^. We found no differences in the proportions of TRPV1-immunoreactive neurons or ATF-3-positive cells in chloroprocaine+capsaicin-treated animals, indicating no ongoing neuronal damage or loss across treatment groups (**Figure 6A**, *upper*, **B**). GFAP expression in the chloroprocaine+vehicle group was similar to that of the saline+vehicle group but higher than on the contralateral side (**Figure 6B**, *lower*, **C**). Taken together, these results indicate that, unlike QX-314-based combinations, chloroprocaine, alone or in combination with capsaicin, does not induce TRPV1-mediated calcium toxicity, delayed behavioral hypersensitivity, or structural neuronal injury, supporting its safety when co-applied with TRPV1 activators.

**Figure 6.**
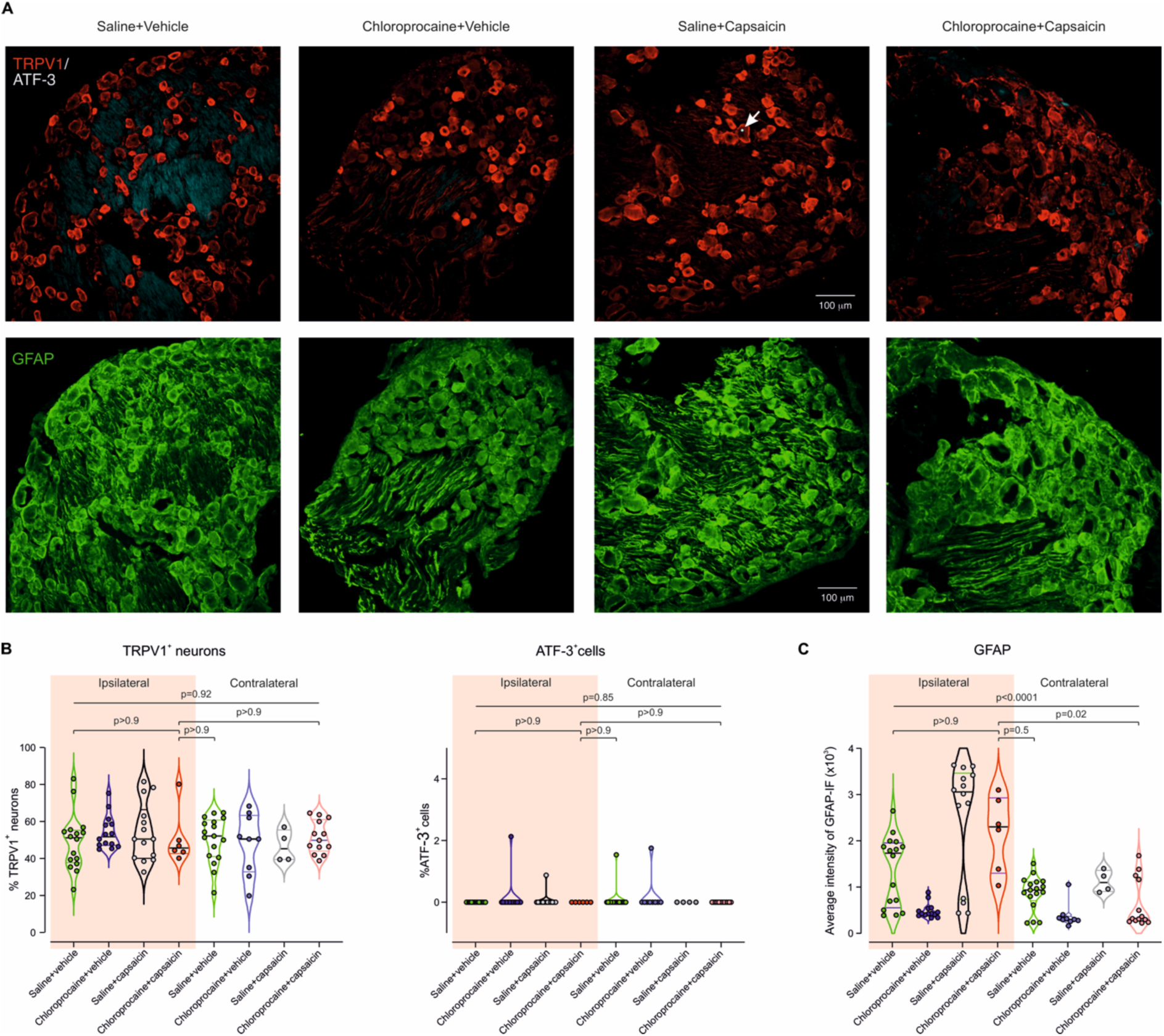
Perisciatic chloroprocaine-capsaicin does not induce neuronal loss or ATF-3 up-regulation but increases DRG GFAP, similar to capsaicin alone. **(A)** Representative confocal images of ipsilateral L4-L6 DRG 35 days after perisciatic injection of saline+vehicle, chloroprocaine+vehicle, saline+capsaicin, or chloroprocaine+capsaicin, showing TRPV1/ATF-3 double labeling (*top*) and GFAP immunoreactivity (*bottom*). Scale bar, 100 μm. The white arrow indicates a cell soma positively stained for ATF-3. (**B**) Comparison of the percentage of TRPV1-positive neurons (*left*) and ATF-3-positive neurons (*right*) among total DAPI-positive neurons following the treatments indicated below, per section in DRGs, ipsilateral and contralateral to the treatment application. (**C**) Same as *B*, but comparison of mean GFAP fluorescence intensity. Each symbol represents one section; violin plots depict the distribution and median. Because sections from different animals were pooled, analyses were performed at the section level using two-way linear models with factors “treatment” and “side” (ipsilateral vs contralateral) and their interaction; post hoc pairwise comparisons were adjusted with Tukey’s test. N = 3 rats per treatment.

### Co-application of chloroprocaine with cannabidiol produces pain-selective analgesia

Our data show that activation of TRPV1 channels by capsaicin can facilitate the entry of chloroprocaine and create a differential block; however, we and others have demonstrated that capsaicin application is potentially neurotoxic (^34^; see also **Figure 6B**). Furthermore, it induces an “injection pain”^17^. The potential toxicity and possible irritative effect of capsaicin could limit the clinical application of a chloroprocaine+capsaicin combination. Therefore, we examined whether we could substitute capsaicin with a less painful TRPV1 agonist^35,36^. It has been demonstrated that the phytocannabinoid cannabidiol (CBD) activates TRPV1 channels. We found that replacing capsaicin with CBD preserved the capacity for pain-selective anesthesia (**Figure 7**). Perisciatic co-administration of chloroprocaine+CBD has an effect similar to that of chloroprocaine+capsaicin (AUC_Chloroprocaine+CBD_ = 9.7±1.8, N=6 rats, p = 0.32, compared with AUC_Chloroprocaine+capsaicin_, Welch’s two-tailed t-test; **Figure 7B**). We and others demonstrated that CBD inhibits nociceptor sodium channels^37–39^, and therefore could potentially induce an analgesic effect when applied alone. However, we show that application of CBD did not produce any analgesic effects when applied together with saline (**Figure 7B**). But when co-applied with chloroprocaine significantly increased thermal withdrawal latency compared with chloroprocaine+vehicle (**Figure 7B**). This decreased sensitivity to noxious heat lasted about 4 hours, while chloroprocaine+CBD induced motor impairment persisted only 30 min and was indistinguishable from chloroprocaine+vehicle (**Figure 7A**). Considering that no pain or hyperalgesia were reported following injection of CBD^40^, a co-application of CBD with chloroprocaine could produce painless pain-selective anesthesia.

**Figure 7.**
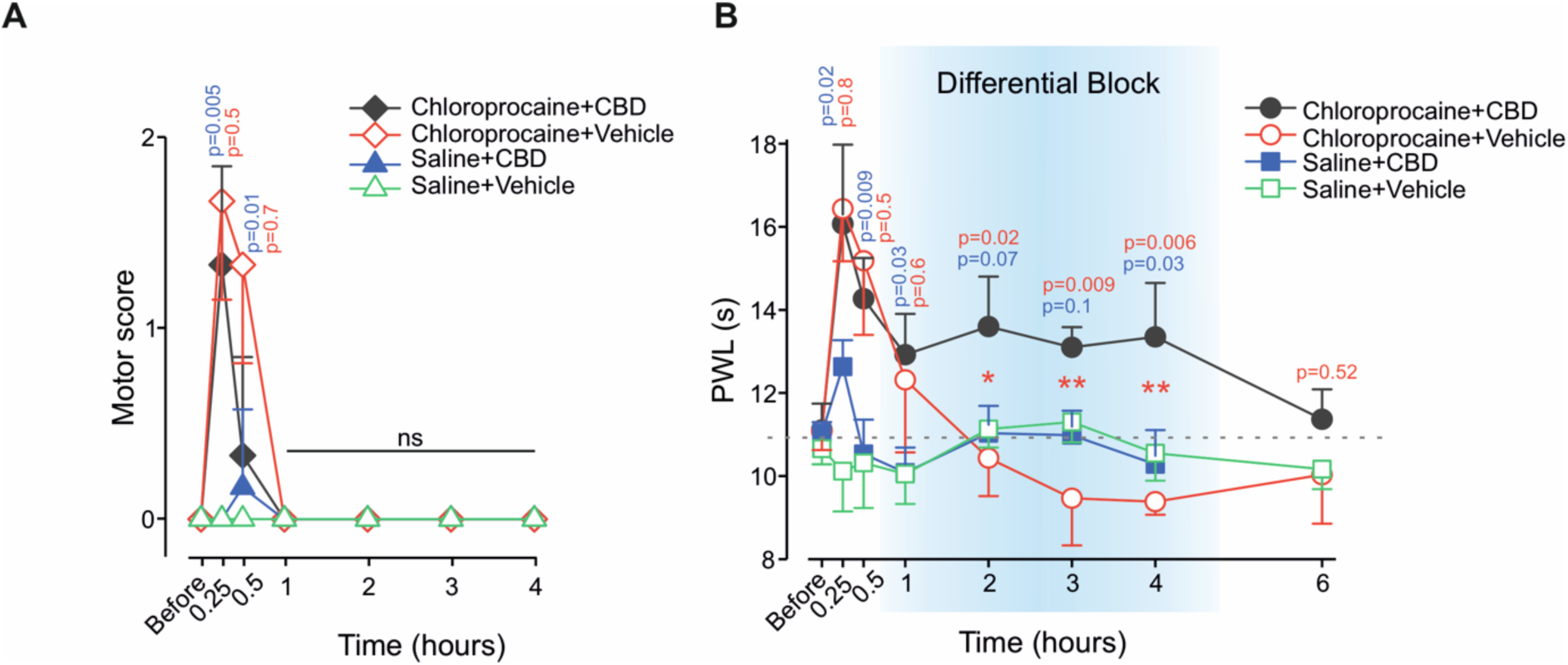
Cannabidiol substitutes for capsaicin to produce TRPV1-mediated pain-selective chloroprocaine block. **(A)** Motor scores over time after perisciatic injection of 0.5% chloroprocaine followed by cannabidiol (CBD, chloroprocaine+CBD), 0.5% chloroprocaine followed by vehicle (chloroprocaine+vehicle), saline followed by CBD (saline+CBD), or saline followed by vehicle (saline+vehicle). (**B**) Changes in sensitivity to noxious thermal stimuli, measured as thermal paw-withdrawal latency (PWL) in the same animals, showing a prolonged increase in latency after chloroprocaine+CBD compared with controls. Shaded regions indicate the period of differential block, during which nociceptive function remained suppressed after the motor block resolved. The dotted lines indicate the mean value before treatment. Data are presented as mean ± SEM. Motor scores were analyzed using generalized estimating equations with an ordinal cumulative logit link, followed by post hoc Mann-Whitney tests with Holm correction at each time point. Changes in PWL were analyzed by two-way repeated-measures ANOVA followed by Tukey’s multiple-comparisons test. N = 6 rats per group. Statistical comparisons between the chloroprocaine+CBD and chloroprocaine+vehicle groups are indicated in *red*. Because of the potential analgesic effect of CBD, we also compared the chloroprocaine+CBD group with the saline+CBD group (*blue*). Exact p-values are shown on the graph.

## Discussion

In this study, we demonstrate that LA with a high pKa, chloroprocaine, produces prolonged, nociceptor-selective anesthesia when co-administered with TRPV1 agonists. This approach is conceptually based on the previously described method for pain-selective anesthesia, which uses the permanently charged lidocaine derivative QX-314. When applied together with the TRPV1 agonist capsaicin, otherwise membrane-impermeant QX-314 reaches the intracellular space of TRPV1-expressing neurons, leading to a modality-specific blockade^6,13^. However, QX-314 causes long-lasting neurotoxicity in part due to TRPV1 channel activation^18,21^. Here, using the clinically approved chloroprocaine, our approach achieves differential blockade without neurotoxicity.

Our approach is grounded in the physicochemical properties of chloroprocaine. With a pKa of 9.1, chloroprocaine is >98% ionized at physiological pH, existing predominantly in its protonated, charged form, which limits lipid membrane permeability relative to other LAs. We hypothesized that opening large-pore TRPV1 channels on nociceptors would provide an alternative entry pathway for the charged fraction, bypassing passive membrane diffusion (**Figure 8**). Our electrophysiological data support this hypothesis: co-application of chloroprocaine with capsaicin produced significantly greater sodium current blockade than either agent alone, indicating TRPV1-facilitated intracellular delivery. This enhanced blockade translated into prolonged behavioral analgesia, validating the concept that TRPV1 channels can serve as selective conduits for otherwise membrane-impermeant LA.

**Figure 8.**
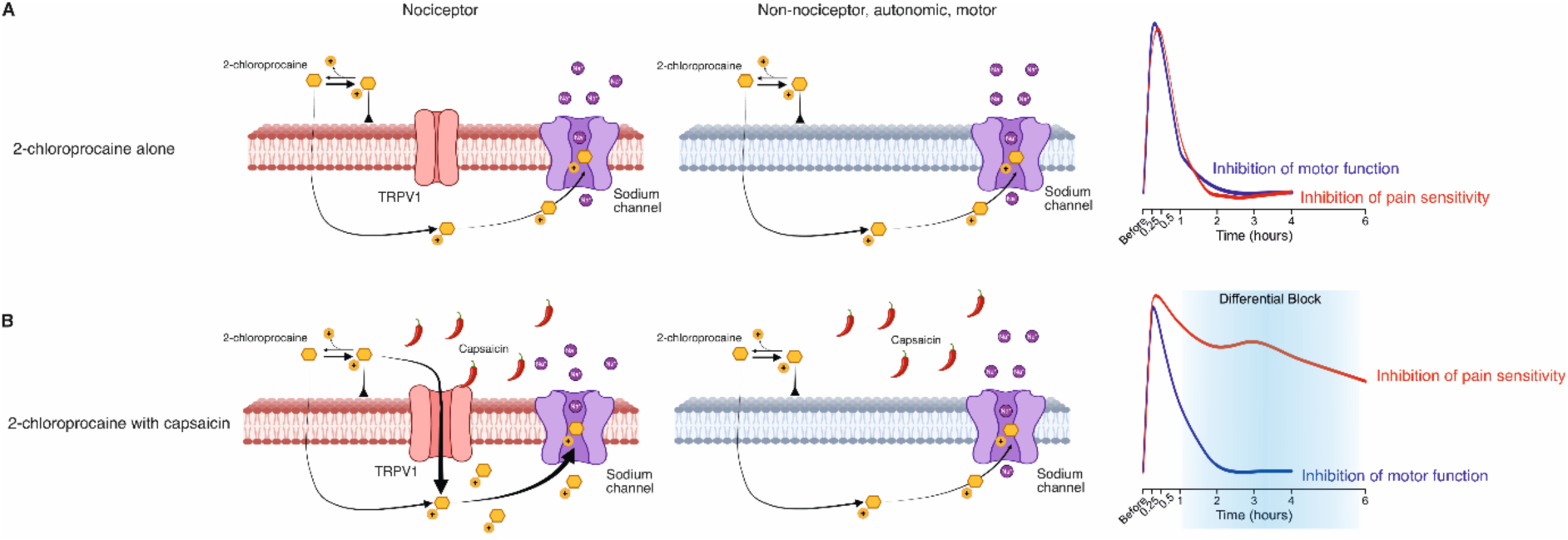
Schematic illustration depicting the differential penetration and action of chloroprocaine in nociceptor versus non-nociceptor neurons. (**A**) The high pKa (9.1) of chloroprocaine favors a predominantly protonated (charged) form at physiological pH, limiting membrane penetration into the cytoplasmic site of sensory and motor neurons, resulting in minimal blockade and rapid recovery of sensory (*red line*) and motor (*blue line*) function. No differential block is produced. (**B**) When chloroprocaine is co-applied with capsaicin or other TRPV1 agonists (e.g., CBD), it opens a large pore on TRPV1-expressing nociceptor neurons. This allows the charged fraction of chloroprocaine to enter selectively through the TRPV1 open channel pore, gaining facilitated access to the intracellular compartment of nociceptor neurons, blocking voltage-gated sodium channels, and producing prolonged analgesia (*red line*). In non-nociceptor neurons (motor and non-nociceptive sensory fibers) that lack TRPV1 expression, chloroprocaine penetrates via passive diffusion, as shown in *A*. The prolonged inhibition of pain sensitivity (*red line*) and the early resolution of motor function (*blue line*) create a differential block.

A critical advantage of chloroprocaine over QX-314 is its well-established safety profile in clinical practice. Chloroprocaine has been used for decades in obstetric and ambulatory surgery, exhibiting rapid plasma hydrolysis by pseudocholinesterase, minimal systemic accumulation, and low maternal-fetal toxicity^41,42^. Clinical studies demonstrate that spinal chloroprocaine provides effective short-duration anesthesia with no reported cases of urinary retention, or persistent neuropathy, complications observed with other LAs^43,44^. Our preclinical findings corroborate this safety record: 24-hour exposure to 1 mM chloroprocaine produced no cytotoxicity in hTRPV1-expressing cells, and long-term behavioral assessment revealed no delayed mechanical hypersensitivity or thermal hyperalgesia up to 35 days post-injection.

The concept of combining capsaicin with LAs is not novel. Previously, we co-applied capsaicin with the tertiary amine LAs lidocaine and bupivacaine, and with amitriptyline and showed that each produced predominantly nociceptive-specific blockade^45^. Critically, unlike lidocaine and other tertiary amine LAs that activate TRPV1 channels^17,46^, we found that chloroprocaine does not activate TRPV1 even at supraclinical concentrations (100 mM). These results are supported by a clinical observation showing that chloroprocaine injection is less painful than lidocaine^47^. Because lidocaine-induced activation of TRPV1 channels has been implicated in injection pain^46^, the lack of TRPV1 activation by chloroprocaine likely accounts for its reduced injection-associated pain. This lack of TRPV1 activation is also mechanistically significant: QX-314 neurotoxicity is mediated by excessive calcium influx through TRPV1 channels it activates^20,21^, leading to calcium-dependent cell death. By not activating TRPV1, chloroprocaine avoids this toxicity mechanism. Our immunohistochemical analysis confirmed the absence of neuronal injury markers (ATF-3) and the preservation of TRPV1-expressing neuron populations after chloroprocaine+capsaicin treatment, contrary to the documented neurotoxicity of QX-314-based combinations^18^. The modest GFAP upregulation observed in our study was attributable to capsaicin alone, not to chloroprocaine or the combination, consistent with previous reports of capsaicin-induced glial activation^48^.

While QX-314 produces pain-specific anesthesia without motor blockade^6^, we observed that 0.5% chloroprocaine+capsaicin produced transient motor impairment lasting 30 minutes, followed by prolonged thermal analgesia persisting beyond 4 hours. This pattern constitutes pain-selective (differential block) rather than pain-specific anesthesia. The brief motor blockade likely reflects some degree of passive membrane penetration by the unionized fraction of chloroprocaine (∼2% at pH 7.4), sufficient to transiently affect motor fibers but insufficient for prolonged blockade. Rapid recovery of motor function occurs as chloroprocaine is hydrolyzed systemically, while nociceptors remain blocked due to the continued intracellular presence of chloroprocaine delivered via TRPV1 channels. This differential blockade profile may offer clinical advantages. Complete intraoperative blockade (sensory and motor) is often desirable during surgery, with rapid motor recovery and sustained postoperative analgesia. Our approach provides precisely this temporal pattern: initial comprehensive blockade transitioning to isolated pain relief as motor function returns. This differs from conventional long-acting LAs (e.g., bupivacaine), which produce prolonged motor blockade that can delay ambulation and hospital discharge.

We observed a more pronounced blockade of thermal pain than of mechanical pain. TRPV1-expressing nociceptors are polymodal but show preferential sensitivity to noxious heat (>42°C) and chemical stimuli^49,50^. Genetic ablation studies demonstrate that TRPV1-expressing neurons are essential for thermal nociception but contribute less prominently to mechanical pain^51^. Therefore, preferential blockade of thermal pain with TRPV1-targeted delivery is mechanistically consistent with the biology of TRPV1-expressing nociceptor subpopulations. This modality selectivity was less apparent in previous QX-314 studies^6^, potentially due to differences in experimental design, drug concentrations, or the contribution of TRPV1 activation by QX-314 itself ^21^. The clinical relevance of modality-specific analgesia warrants investigation, as different surgical procedures and pain conditions may benefit differentially from thermal versus mechanical pain blockade.

We found that a 10-minute interval between perineural chloroprocaine and capsaicin injections optimized the differential block, whereas a 2-minute interval was more effective for intraplantar injection. This temporal requirement likely reflects the kinetics of LA tissue penetration, systemic absorption, and TRPV1 channel availability. We propose that the optimal interval allows extracellular accumulation of chloroprocaine while TRPV1 channels are activated, thereby maximizing the availability of both the drug and the entry pathway. Local tissue pH changes caused by the acidic formulations of both agents might transiently alter TRPV1 activation kinetics or LA ionization states. The shorter optimal interval for intraplantar injection likely reflects the smaller tissue volume and more rapid drug equilibration in paw tissue compared with the perisciatic space. These pharmacokinetic considerations will be important for optimizing clinical protocols and may require titration based on injection site, volume, and desired duration of effect.

Although capsaicin effectively facilitated chloroprocaine entry and produced robust differential blockade, it has limitations: it causes acute “injection pain” and induces glial activation, as observed in our GFAP immunohistochemistry. We therefore explored cannabidiol (CBD), a non-psychotropic phytocannabinoid that activates TRPV1 channels through distinct mechanisms, including channel desensitization and modulation of adenylyl cyclase-cAMP signaling^52,53^. CBD-mediated TRPV1 activation is milder than capsaicin, producing calcium influx without the intense pain sensation^54^. Our results demonstrate that substituting capsaicin with CBD preserved pain-selective analgesia comparable to capsaicin. Thus, chloroprocaine+CBD combinations may be more clinically acceptable than capsaicin-based formulations. Although CBD can directly inhibit sodium channels at high concentrations^37–39^, we observed no analgesic effect of CBD alone (saline+CBD group), indicating that the enhanced effect of chloroprocaine+CBD results from TRPV1-mediated facilitation rather than direct channel blockade by CBD.

Another potential substitute for capsaicin is lidocaine. It has been shown that lidocaine activates TRPV1 channels^46^. Moreover, we demonstrated that lidocaine-induced activation of TRPV1 channels is sufficient to allow QX-314 entry into the nociceptor intracellular space and to produce differential blockade^17,55^. However, recently it has been shown that combining two LAs significantly shortened the duration of analgesia without affecting the sensory onset time^56^. This could result from complex interactions between two membrane-permeant LAs, which may alter tissue pharmacokinetics^56^. Thus, combining lidocaine to activate TRPV1 channels with chloroprocaine, although a logically sound approach, is not trivial and may trigger complex interactions with unexpected results.

The key innovation of our approach is combining two clinically used agents - chloroprocaine (FDA-approved LA) and potentially CBD (increasingly available for medical use) - to achieve a therapeutic effect (pain-selective anesthesia) not attainable with either agent alone. The established safety profiles of both chloroprocaine and CBD may facilitate clinical translation. However, optimal concentrations, volumes, and ratios of chloroprocaine to TRPV1 agonists require systematic dose-finding studies. Safety assessment in humans, particularly long-term neurological follow-up and assessment of injection site reactions, will be essential before widespread clinical adoption.

Although our data are consistent with TRPV1-mediated entry of chloroprocaine, we cannot exclude indirect mechanisms such as capsaicin-induced depolarization, which activates and subsequently inactivates sodium channels, “sensitizing” them to LA blockade by virtue of their higher affinity for inactivated channels^57^. Also, TRPV1 activation acidifies the cytoplasm^58^, increasing the proportion of LA in the charged form intracellularly, which is generally more potent than the neutral form and leaves the cell more slowly ^59,60^. Finally, calcium entry through TRPV1 channels may alter sodium channel phosphorylation states, potentially enhancing LA efficacy.

In conclusion, we demonstrate that the clinically approved high-pKa local anesthetic chloroprocaine, when co-administered with TRPV1 agonists such as capsaicin or cannabidiol, produces prolonged pain-selective analgesia without evidence of neurotoxicity. This effect is likely mediated by TRPV1-facilitated entry of the predominantly protonated, membrane-impermeant fraction of chloroprocaine into nociceptive neurons, resulting in sustained inhibition of voltage-gated sodium channels. Importantly, chloroprocaine does not activate TRPV1 channels, thereby avoiding the calcium-dependent toxicity mechanisms implicated in prior TRPV1-based approaches. Together, these findings establish a potentially translatable strategy for pain-selective regional anesthesia that leverages the physicochemical properties of clinically approved high-pKa local anesthetics. This approach may have broad applications in perioperative pain management, labor analgesia, and chronic pain treatment. Further clinical studies will be required to optimize formulations, dosing, and long-term safety in human patients.

## Ethical approval

This study was approved by the Hebrew University IACUC under Approval numbers MD-18-15551-5 and MD-24-17427-5.

## Previous presentation

partial initial findings from this study were presented in abstract form in the World Conference of Veterinary Anesthesia in Venice 27^th^ September 2018, and the World Conference of Veterinary Anesthesia in Paris 16^th^ September 2025.

## Acknowledgements

**Summary statement (35 words):** This animal study demonstrates that nociceptive selective blockade without neurotoxicity can be achieved by combining a clinically utilized TRPV1 agonist and a polarized local anesthetic, 2-chloroprocaine, thus providing loco-regional anesthesia without motor blockade.

## Supplementary Figures

**Figure S1.**
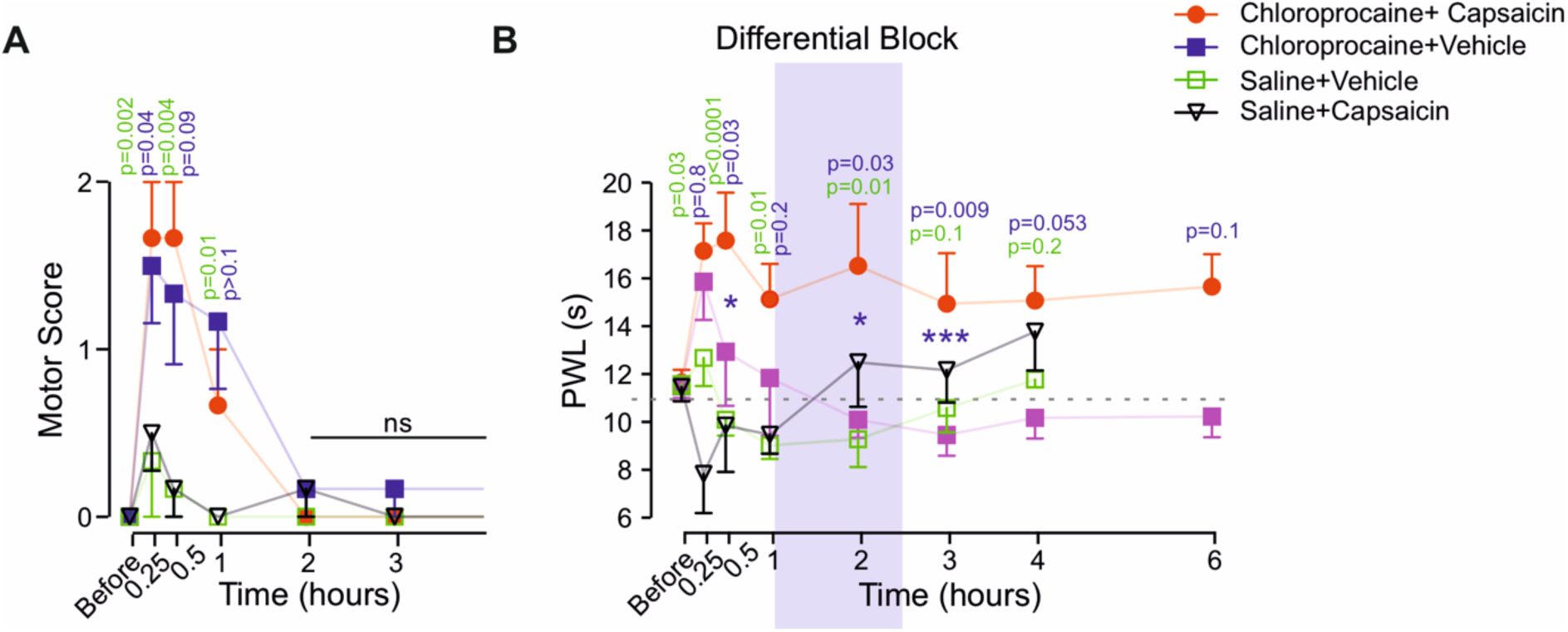
Higher dose (2%) chloroprocaine co-applied with capsaicin does not improve differential block. **(A)** Changes in motor scores (0-2) over time after perisciatic injection of 2% chloroprocaine followed by 0.05% capsaicin (chloroprocaine+capsaicin), 2% chloroprocaine followed by vehicle (chloroprocaine+vehicle), saline followed by 0.05% capsaicin (saline+capsaicin), or saline followed by vehicle (saline+vehicle). (**B**) Changes in the sensitivity to noxious thermal stimuli (paw withdrawal latency, PWL) over time following the same treatments as in *A*. Shaded regions indicate the period of differential block, during which nociceptive function remained suppressed after the motor block resolved. The dotted lines indicate the mean value before treatment. Data are mean ± SEM. Motor scores were analyzed using generalized estimating equations; between-group differences at individual time points were further assessed with Mann-Whitney tests with Holm-Bonferroni correction. PWL were analyzed with two-way repeated-measures ANOVA followed by Tukey’s multiple-comparisons test. N = 6 rats per group. Statistical comparisons between the chloroprocaine+capsaicin and chloroprocaine+vehicle groups are indicated in blue, and between chloroprocaine+capsaicin and saline+vehicle in green. Exact p-values are shown on the graph.

**Figure S2.**
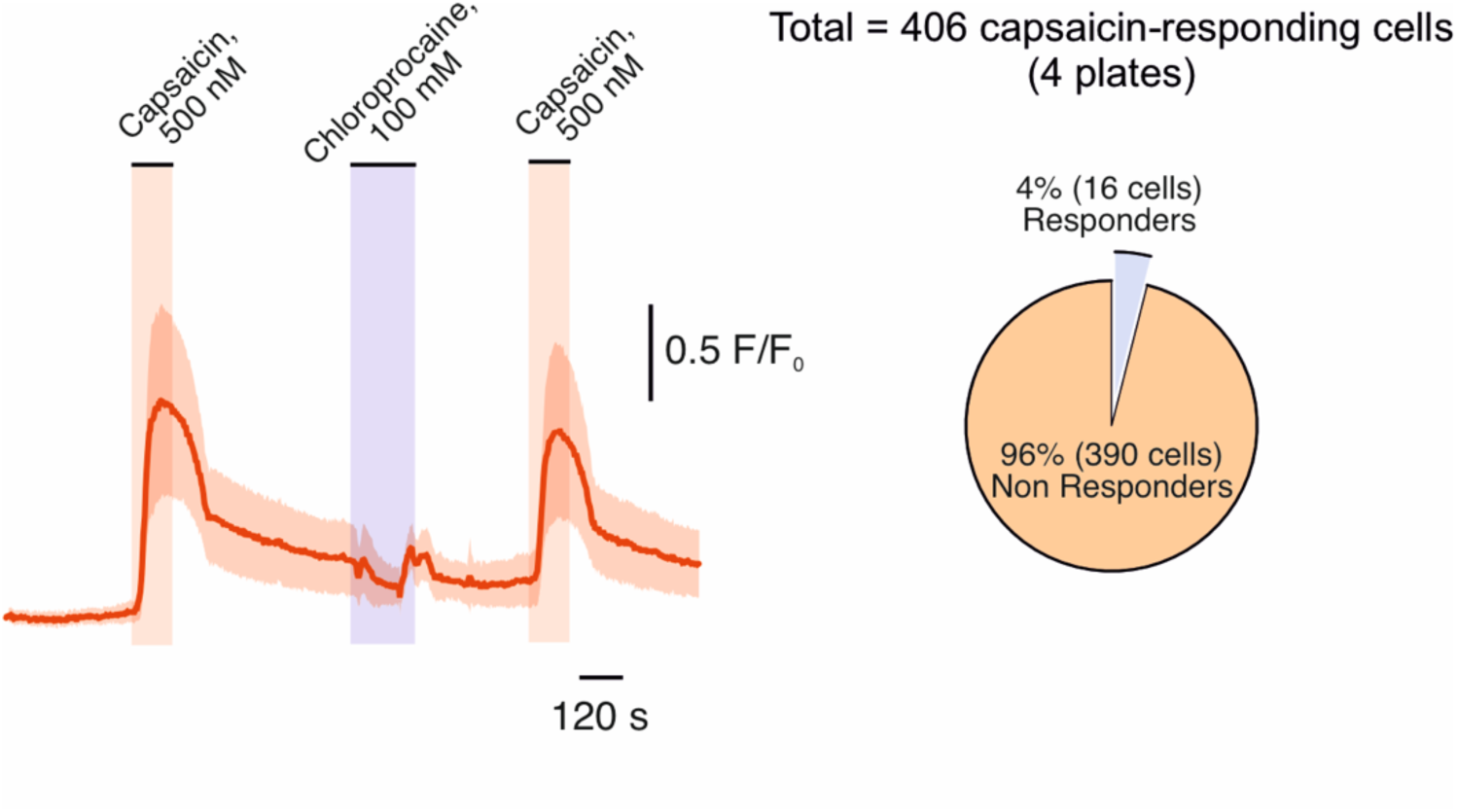
High concentrations of chloroprocaine (100 mM) do not activate TRPV1 in HEK293-hTRPV1 cells. Representative trace of mean changes in Ca^2+^ responses (measured as ratiometric changes in Fura-2 fluorescence; the cloud shows the full range of responses) in HEK293 cells expressing hTRPV1 following application of 100 mM chloroprocaine and 500 nM capsaicin. The abrupt bidirectional changes after the application of 100 mM chloroprocaine are a non-specific response due to changes in absorption at 380 nm and not at 340 nm, possibly due to the high osmolarity of the solution. The pie chart shows the proportion of capsaicin-responsive cells that responded to chloroprocaine. Representative of 4 out of 4 plates.

