## Supplemental Figures 1, 2 for "TRPV1-Mediated Delivery of Chloroprocaine, a Local Anesthetic with High pKa, Produces Pain-Selective Anesthesia Without Neurotoxicity"

### Supplementary Figures

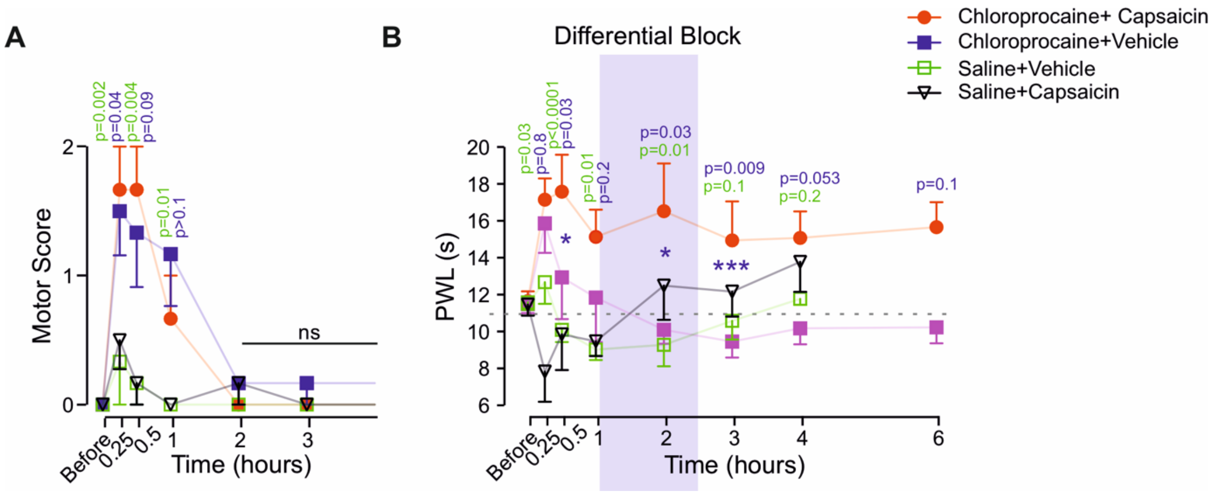


**Figure S1. Higher dose (2%) chloroprocaine co-applied with capsaicin does not improve differential block. (A)** Changes in motor scores (0-2) over time after perisciatic injection of 2% chloroprocaine followed by 0.05% capsaicin (chloroprocaine+capsaicin), 2% chloroprocaine followed by vehicle (chloroprocaine+vehicle), saline followed by 0.05% capsaicin (saline+capsaicin), or saline followed by vehicle (saline+vehicle). (**B**) Changes in the sensitivity to noxious thermal stimuli (paw withdrawal latency, PWL) over time following the same treatments as in *A*. Shaded regions indicate the period of differential block, during which nociceptive function remained suppressed after the motor block resolved. The dotted lines indicate the mean value before treatment. Data are mean ± SEM. Motor scores were analyzed using generalized estimating equations; between-group differences at individual time points were further assessed with Mann-Whitney tests with Holm-Bonferroni correction. PWL were analyzed with two-way repeated-measures ANOVA followed by Tukey's multiple-comparisons test. N = 6 rats per group. Statistical comparisons between the chloroprocaine+capsaicin and chloroprocaine+vehicle groups are indicated in blue, and between chloroprocaine+capsaicin and saline+vehicle in green. Exact p-values are shown on the graph.

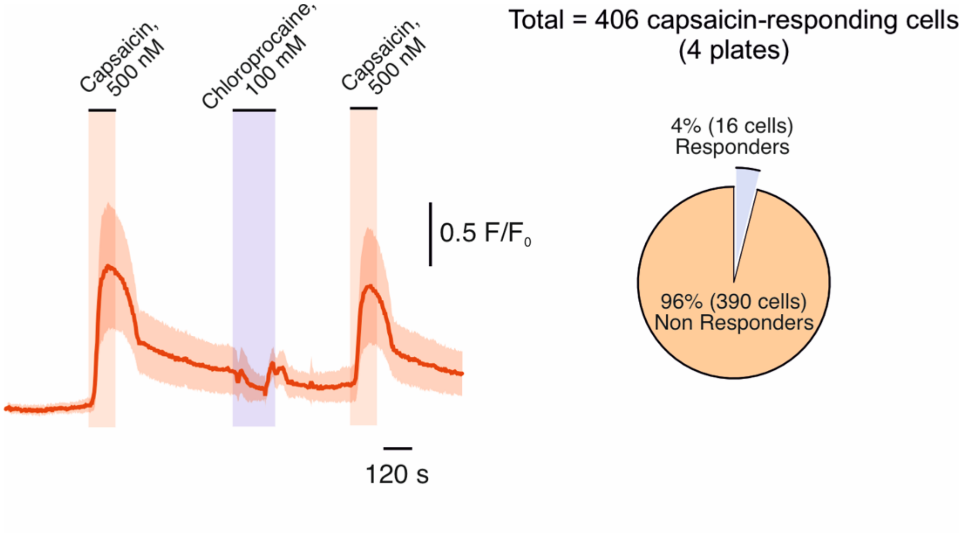


**Figure S2. High concentrations of chloroprocaine (100 mM) do not activate TRPV1 in HEK293-hTRPV1 cells.** Representative trace of mean changes in $Ca^{2+}$ responses (measured as ratiometric changes in Fura-2 fluorescence; the cloud shows the full range of responses) in HEK293 cells expressing hTRPV1 following application of 100 mM chloroprocaine and 500 nM capsaicin. The abrupt bidirectional changes after the application of 100 mM chloroprocaine are a non-specific response due to changes in absorption at 380 nm and not at 340 nm, possibly due to the high osmolarity of the solution.  The pie chart shows the proportion of capsaicin-responsive cells that responded to chloroprocaine. Representative of 4 out of 4 plates.
